# HUMAN FETAL CIRCULATING FACTORS FROM PREGNANCIES COMPLICATED BY MATERNAL OBESITY INDUCE HYPERTROPHY IN NEONATAL RAT CARDIOMYOCYTES

**DOI:** 10.1101/2025.04.16.649071

**Authors:** Owen R. Vaughan, Andrew Goodspeed, Carmen C. Sucharov, Theresa L. Powell, Thomas Jansson

## Abstract

**Aim:** Obesity in pregnant women increases offspring cardiovascular risk and causes fetal cardiac dysfunction. The underpinning mechanisms remain unclear. We hypothesised that circulating factors in serum from fetuses of women with obesity induce pathological cardiomyocyte hypertrophy.

**Methods:** Pregnant women with obesity or healthy weight were recruited at term and provided umbilical cord serum and placentas, which were used for isolation of primary trophoblast cells. Primary cardiomyocytes were isolated from neonatal rats.

**Results:** Compared to serum-free medium, supplementing cardiomyocytes with umbilical cord serum increased their mRNA expression of atrial natriuretic factor (*Anf*) and brain natriuretic peptide (*Bnp*), hallmarks of pathological hypertrophy, but did not alter their size. Cord serum from women with obesity further upregulated cardiomyocyte *Anf* and *Bnp*, and increased the ratio of beta- to alpha-myosin heavy chain expression (*Myh7:Myh6*), compared to cord serum from healthy weight women. This effect was prevented by treating the cord serum with heat-freeze cycling and DNase or RNase digestion. Conditioned medium from trophoblast cells from women with obesity also increased cardiomyocyte *Anf*, *Bnp* and *Myh7:Myh6* expression. MicroRNAs miR-142 and miR-17, which are associated with cardiac function, were increased in abundance in extracellular vesicles isolated from cord serum from women with obesity. However, miR-142-3p, miR-142-5p and miR-17-5p did not increase *Anf*, *Bnp* or *Myh7:Myh6* expression when they were transfected into cardiomyocytes.

**Conclusion:** The results show that human fetal circulating and placenta-derived factors induce hallmarks of pathological hypertrophy in cardiomyocytes and may mediate cardiac dysfunction in children of women with obesity.

## INTRODUCTION

Obesity in pregnant women is associated with increased risk for cardiometabolic disease in the offspring throughout life ^1^. This predisposition may be established *in utero* by adverse alterations in the structure and function of the fetal heart. Maternal obesity increases the thickness of the fetal left ventricular walls and intraventricular septum and reduces cardiac strain rate ^2,3^, consistent with cardiac hypertrophy and contractile dysfunction. Cardiac dysfunction persists postnatally in the offspring, at least into childhood ^4^. Experimentally- induced obesity in pregnant animals similarly causes fetal cardiac dysfunction in association with hypertrophy and impaired contractility of individual cardiomyocytes ^5–10^. The mechanisms that induce fetal cardiomyocyte hypertrophy in pregnancies complicated by obesity remain unknown.

Humoral factors contribute mechanistically to pathological cardiac hypertrophy in postnatal life and catecholamines, angiotensin II, inflammatory cytokines, growth factors and insulin are all implicated ^11^. For example, serum samples from patients with pediatric dilated cardiomyopathy induce hypertrophy in cardiomyocytes *in vitro* in a β-adrenergic-dependent manner ^12^. Maternal obesity alters the fetal hormonal and metabolic milieu, and cord plasma circulating factors are strongly associated with cardiac function in the infant ^13^ .

Recently, circulating nucleic acids including microRNAs have been shown to mediate inter-organ crosstalk when they are stabilised in extracellular vesicles or carrier proteins, and cause cardiomyocyte hypertrophy ^14,15^. MicroRNAs bind to specific target messenger RNAs to regulate their degradation and/or translation and are known to influence intracellular proteins that are important in cardiomyocyte hypertrophy. Obesity in pregnant women alters the maternal circulating and placental abundance of microRNAs that predict the postnatal growth and metabolic phenotype of the infant ^16,17^. Maternal obesity also alters microRNA expression in the fetal heart in non-human primates ^18^. The contribution of circulating factors and microRNAs to fetal cardiac hypertrophy, as opposed to mechanical or neural inputs, remains to be established.

Fetal cardiac development is linked to placental function. The placental circulation and fetal heart develop in parallel in early gestation ^19^. Genetic manipulations specifically targeted to placental trophoblast cells alter fetal cardiac phenotype in mice ^20^. In pregnant mice with diet-induced obesity, maternal adiponectin supplementation normalises both placental function and postnatal cardiac dysfunction ^21^. Placental cells synthesise hormones, metabolites, signalling molecules and extracellular vesicles that are released into the fetal circulation ^22–24^. Placenta-secreted factors may therefore play a role in fetal cardiac development in pregnancies complicated by maternal obesity.

We hypothesised that circulating factors in cord serum from fetuses of women with obesity induce pathological cardiomyocyte hypertrophy. We aimed to determine the effect of fetal circulating factors on cultured primary neonatal rat ventricular cardiomyocytes, a well-established model used to investigate mechanisms of pathological hypertrophy. We also tested the cardiomyocyte effects of circulating microRNAs and placenta-derived factors.

## RESULTS

### Effect of human umbilical cord serum on cardiomyocytes *in vitro*

Umbilical cord serum from infants of women with healthy weight or obesity was added to the medium of primary neonatal rat ventricular myocytes. By definition, women with obesity had significantly higher pre-pregnancy BMI than healthy weight controls but the two groups had similar maternal ethnicity, age, GDM status, gestational age, delivery mode, birth weight, placenta weight and fetal sex (**Table 1**). Cord serum from fetuses of healthy weight control women upregulated NRVM expression of the fetal genes *Anf* and *Bnp*, compared to culturing in serum-free medium (**Fig 1A, B**). Cord serum from fetuses of women with obesity further upregulated *Anf* and *Bnp*, compared either to control serum or serum-free medium (**Fig 1A,B**). The ratio of fetal *Myh7* to adult *Myh6* expression was greater in NRVMs supplemented with cord serum from fetuses of women with obesity, but not healthy weight women, compared to serum-free NRVMs (**Fig. 1C**). *Serca* expression was not affected by cord serum supplementation (**Fig. 1D**). Cardiomyocyte size, determined by α-actinin immunofluorescence, was also similar in NRVMs with or without cord serum (healthy BMI cord serum 1756 ± 209 µm^2^, maternal obesity cord serum 1664 ± 211 µm^2^, no serum 1748 ± 184 µm^2^, P=0.637).

**Figure 1.**
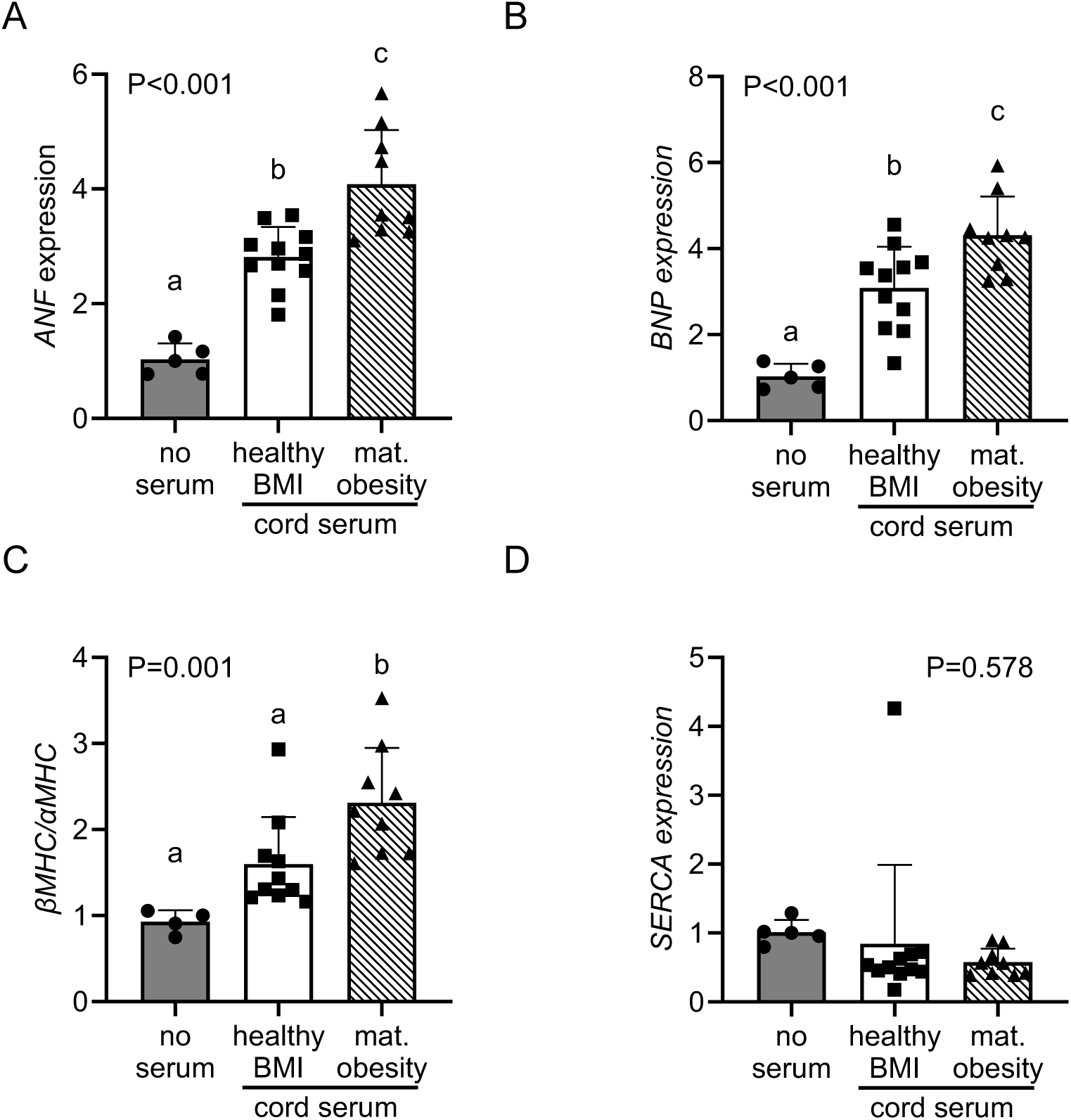
Effect of human umbilical cord serum on cardiomyocytes. (A-D) mRNA expression of fetal gene program markers of pathological hypertrophy in NRVMs in serum-free medium (n=5) or in medium supplemented with cord serum from women with healthy weight (con, n=10) or obesity (n=9). Effect of serum treatment determined by one-way ANOVA. Main effect P value given in figure. Letters a, b, c represent significantly different groups (P<0.05) by Tukey’s post-hoc comparison. Mean ± SD, points represent means of technical replicate cell wells treated with serum from one study participant.

**Table 1.** Clinical characteristics of human study participants providing cord serum for NRVM experiment.

|  | **Healthy weight** | **n** | **Obesity** | **n** | **P value** |
| --- | --- | --- | --- | --- | --- |
| **Pre-pregnancy BMI, kg/m<sup>2</sup>** | 22.6 ± 1.8 | 10 | 37.5 ± 3.2 | 9 | **\*P<0.001** |
| **Ethnicity, number Hispanic (%)** | 1 (10%) | 10 | 4 (44%) | 9 | P=0.089 |
| **Maternal age, years** | 31.7 ± 4.4 | 10 | 29.2 ± 6.3 | 9 | P=0.327 |
| **Gestational diabetes, number (%)** | 1 (10%) | 10 | 0 (0%) | 9 | P=0.330 |
| **Gestational age at delivery, weeks** | 39.2 ± 0.9 | 10 | 39.9 ± 2.2 | 9 | P=0.693 |
| **Vaginal delivery, number (%)** | 1 (10%) | 10 | 0 (0%) | 9 | P=0.330 |
| **Birth weight, kg** | 3.42 ± 0.18 | 9 | 3.78 ± 1.21 | 9 | P=0.382 |
| **Placenta weight, g** | 653 ± 125 | 10 | 774 ± 287 | 9 | P=0.242 |
| **Sex, number female (%)** | 5 (50%) | 10 | 3 (33%) | 9 | P=0.463 |

Continuous variables compared by Student's t-test. Categorical variables compared by Chi-squared test. Mean ± SD.

Exposing cord serum to repeated heat-freeze cycles before supplementing it into the NRVM medium tended to prevent the effect of maternal obesity, such that cardiomyocyte *Anf* expression was no longer upregulated relative to control serum treatment (**Fig. 2A**). Serum heat- freezing also significantly reduced the cardiomyocyte *Myh7:Myh6* ratio, irrespective of the BMI group providing the serum (**Fig. 2B**). The hypertrophic effect of cord serum from pregnancies complicated by obesity was restored when liposomal transfection reagent was included in the medium after heat-freezing (**Fig. 2C, D**). Treating cord serum with DNase, RNase, or a combination of both before supplementing it into the NRVM medium prevented its effect on *Anf*, such that expression was similar in cells treated with cord serum from healthy weight women or women with obesity (**Fig. 2C**). Serum nuclease treatment also reduced the cardiomyocyte *Myh7:Myh6* ratio overall and tended to prevent the effect of cord serum from women with obesity (**Fig. 2D**).

**Figure 2.**
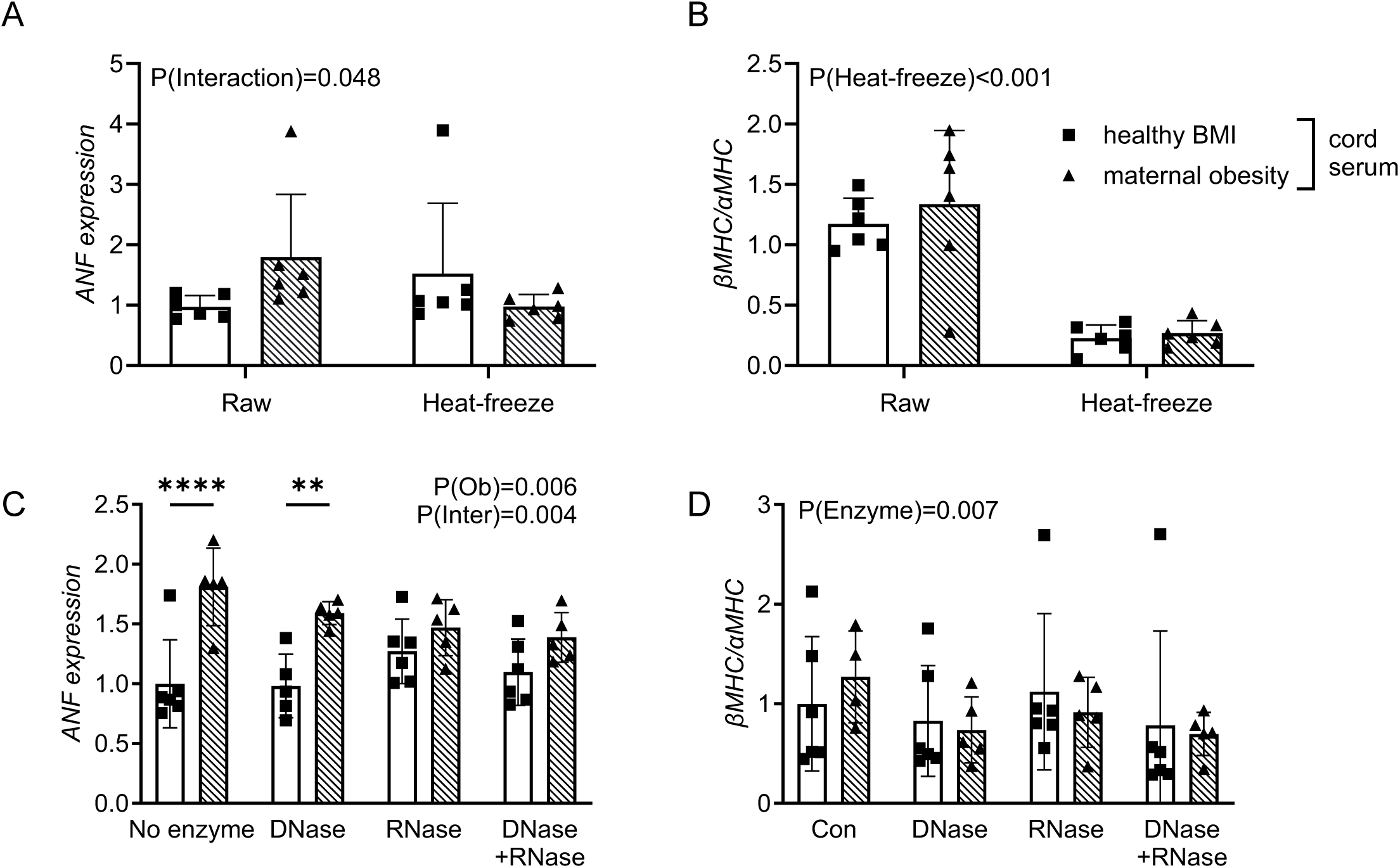
Effect of depleting nucleic acids from cord serum on cardiomyocytes. (A-F) mRNA expression of fetal gene program markers of pathological hypertrophy in NRVMs supplemented with cord serum from women with healthy weight (con, n=6) or obesity (ob, n=5). Effects of maternal obesity (P(Ob)), serum heat-freezing or nuclease treatment (P(Heat-freeze), P(Enzyme)) and interaction (P(Inter) determined by mixed effects model. P values for significant (P<0.05) main effects given in figure. * represent significantly different pairs (P<0.05) by Sidak’s post-hoc comparison. Mean ± SD, points represent means of technical replicate cell wells treated with serum from one study participant.

### Effect of maternal obesity on cord serum microRNA abundance

RNA was purified from extracellular vesicles isolated from cord serum from fetuses of women with healthy weight or obesity. In this larger set of participants, women with obesity had significantly higher pre-pregnancy BMI, heavier babies and greater placental weights than healthy weight controls (**Table 2**). The women with obesity were also more commonly of Hispanic ethnicity but none of the other clinical characteristics differed in the two groups (**Table 2**). Small RNASeq identified 27 upregulated miRNAs and 22 downregulated miRNAs with uncorrected P<0.05 in cord serum from women with obesity, compared to women with healthy weight (**Fig. 3A, Table 3**). The upregulated microRNAs included miR-142-3p, miR-142-5p and miR-17-5p, which are conserved between humans and rodents and have been shown to regulate cardiac function ^25,26^. The experimentally-confirmed downstream mRNA targets of miR- 142-3p, miR-142-5p and miR-17-5p were significantly enriched in transcripts involved in regulating cardiac muscle hypertrophy, metabolic process and growth (**Table 4**). Following multiplicity correction using false discovery rate, maternal obesity significantly upregulated only miR-142-3p and downregulated miR-5010-5p, miR-100-5p, miR483-5p, and miR-149-3p (FDR<0.05, **Fig. 3A, Table 3**).

**Figure 3.**
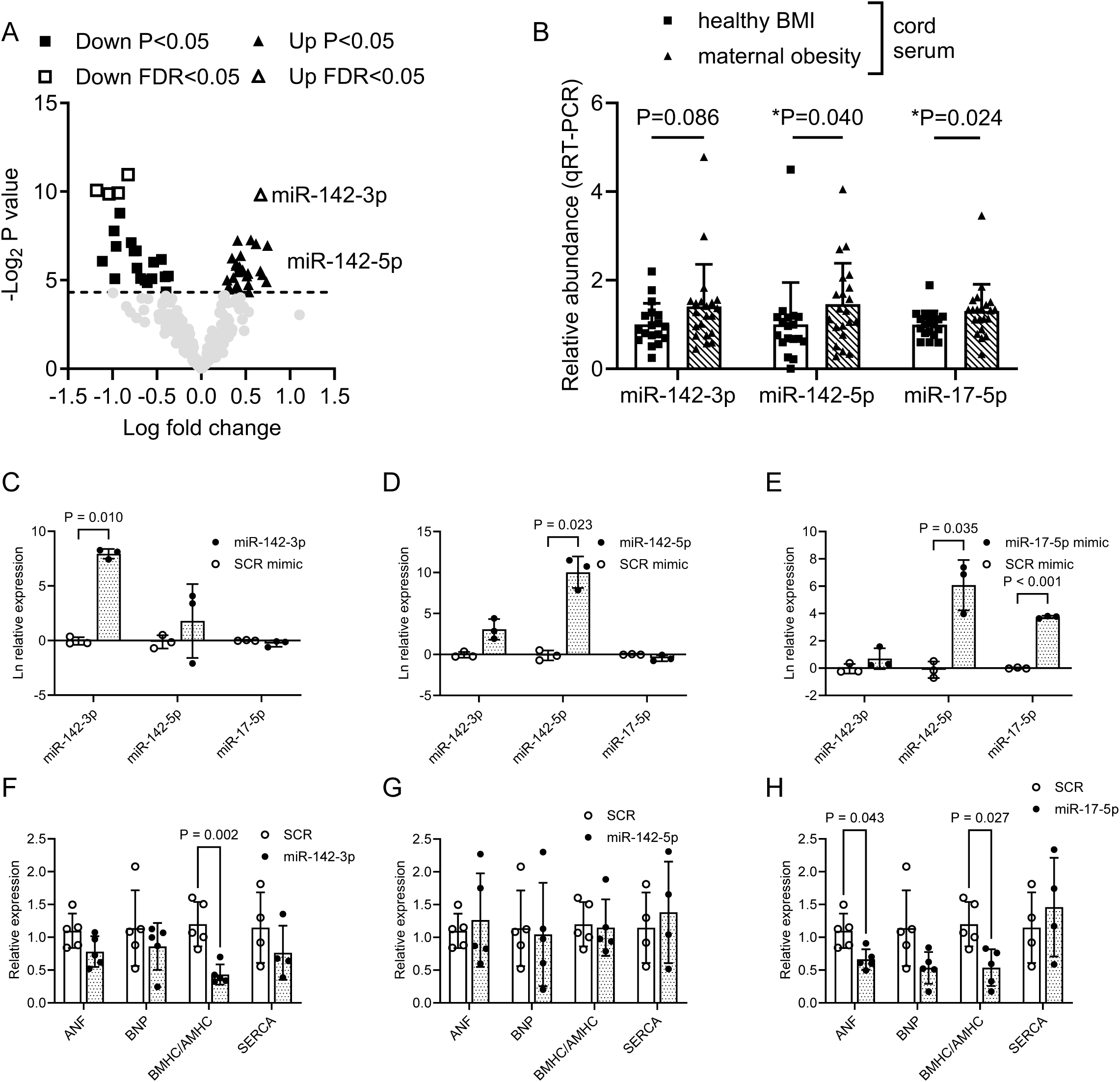
Effect of maternal obesity on fetal circulating microRNA abundance. (A) Volcano plot showing upregulated and downregulated microRNAs by small RNASeq in cord serum from women with obesity (n=14), compared to healthy weight controls (n=12). Points represent individual microRNAs. (B) Relative abundance of selected microRNAs in cord serum from a validation set of women with obesity (n=21) and healthy weight controls (n=18), determined by qT-PCR. Effect of obesity was determined by Student’s t-test. Abundance values were log transformed before statistical analysis. Mean ± SD, points represent individual study participants. (C-E) MicroRNA expression and (F-H) mRNA expression of fetal gene program markers of pathological hypertrophy in NRVMs transfected with synthetic mimics for miR-142- 3p, miR-142-5p or miR-17-5p. n=3-5 independent transfections. Effect of transfection determined by paired Student’s t-test. Mean ± SD, points represent biological replicate experiments in independently isolated batches of primary cardiomyocytes.

**Table 2.** Clinical characteristics of human study participants providing cord serum for RNASeq.

|  | **Healthy weight** | **n** | **Obesity** | **n** | **P value** |
| --- | --- | --- | --- | --- | --- |
| **Pre-pregnancy BMI, kg/m<sup>2</sup>** | 22.6 ± 1.5 | 12 | 37.2 ± 2.9 | 14 | **\*P<0.001** |
| **Ethnicity, number Hispanic (%)** | 1 (8%) | 12 | 7 (50%) | 14 | **\*P=0.022** |
| **Maternal age, years** | 30.3 ± 5.0 | 12 | 30.1 ± 5.6 | 14 | P=0.928 |
| **Gestational age at delivery, weeks** | 39.0 ± 1.05 | 12 | 38.9 ± 1.7 | 14 | P=0.911 |
| **Gestational diabetes, number (%)** | 2 (17%) | 12 | 2 (14%) | 14 | P=0.867 |
| **Vaginal delivery, number (%)** | 2 (17%) | 12 | 1 (7%) | 14 | P=0.449 |
| **Birth weight, kg** | 3.22 ± 0.36 | 11 | 3.94 ± 1.06 | 14 | **\*P=0.031** |
| **Placenta weight, g** | 624 ± 136 | 11 | 797 ± 249 | 14 | **\*P=0.050** |
| **Sex, number female (%)** | 8 (67%) | 12 | 4 (29%) | 14 | P=0.052 |

Continuous variables compared by Student's t-test. Categorical variables compared by Chi-squared test. Welch's correction was used if variances differed between groups. Mean ± SD.

**Table 3.** Differentially abundant microRNAs in cord serum from fetuses of women with obesity versus healthy weight, identified by RNASeq.

| microRNA | Fold change (maternal obesity / healthy weight) | P-value | False discovery rate |
| --- | --- | --- | --- |
| ***Upregulated*** |  |  |  |
| i | 1.588 | 0.001 | 0.045 |
| hsa-miR-20a-5p | 1.474 | 0.006 | 0.130 |
| hsa-let-7g-5p | 1.326 | 0.007 | 0.130 |
| hsa-miR-454-3p | 1.534 | 0.008 | 0.130 |
| hsa-miR-301a-3p | 1.678 | 0.008 | 0.130 |
| hsa-miR-17-5p | 1.360 | 0.012 | 0.152 |
| hsa-let-7i-5p | 1.271 | 0.013 | 0.156 |
| hsa-miR-93-5p | 1.313 | 0.016 | 0.159 |
| hsa-miR-30e-3p | 1.311 | 0.018 | 0.162 |
| hsa-miR-26b-5p | 1.349 | 0.019 | 0.162 |
| hsa-miR-142-5p | 1.381 | 0.021 | 0.162 |
| hsa-miR-625-5p | 1.587 | 0.022 | 0.162 |
| ***Downregulated*** |  |  |  |
| hsa-miR-149-3p | 0.565 | 0.001 | 0.045 |
| hsa-miR-5010-5p | 0.442 | 0.001 | 0.045 |
| hsa-miR-483-5p | 0.523 | 0.001 | 0.045 |
| hsa-miR-100-5p | 0.487 | 0.001 | 0.045 |
| hsa-miR-99a-5p | 0.530 | 0.002 | 0.077 |
| hsa-miR-483-3p | 0.506 | 0.005 | 0.130 |
| hsa-miR-10400-5p | 0.579 | 0.007 | 0.130 |
| hsa-miR-3168 | 0.514 | 0.008 | 0.130 |
| hsa-miR-4466 | 0.602 | 0.010 | 0.136 |
| hsa-miR-6858-5p | 0.592 | 0.010 | 0.136 |
| hsa-miR-127-3p | 0.733 | 0.014 | 0.156 |
| hsa-miR-516a-5p | 0.462 | 0.015 | 0.156 |

**Table 4.** Top biological process GO terms significantly enriched with targets of serum miRNAs upregulated by maternal obesity.

| Pathway | Enrichment | P-value |
| --- | --- | --- |
| ***miR-142-3p*** |  |  |
| cellular protein modification process | 1.43 | 1.0E-02 |
| protein modification process | 1.43 | 1.0E-02 |
| enzyme binding | 1.61 | 1.0E-02 |
| positive regulation of metabolic process | 1.45 | 1.4E-02 |
| regulation of protein metabolic process | 1.53 | 1.5E-02 |
| innate immune response-activating signal transduction | 2.89 | 1.7E-02 |
| activation of innate immune response | 2.68 | 1.9E-02 |
| cardiac muscle hypertrophy | 4.75 | 1.9E-02 |
| positive regulation of cardiac muscle hypertrophy | 8.25 | 1.9E-02 |
| positive regulation of immune response | 2.05 | 1.9E-02 |
| ***miR-142-5p*** |  |  |
| negative regulation of growth | 13.76 | 4.6E-04 |
| positive regulation of cell differentiation | 5.18 | 4.6E-04 |
| regulation of transforming growth factor beta2 production | 103.45 | 4.6E-04 |
| cellular response to transforming growth factor beta stimulus | 10.43 | 6.7E-04 |
| enzyme linked receptor protein signaling pathway | 4.83 | 6.7E-04 |
| epithelial to mesenchymal transition | 14.71 | 6.7E-04 |
| negative regulation of cell differentiation | 5.76 | 6.7E-04 |
| negative regulation of oxidative stress-induced intrinsic apoptotic signaling pathway | 69.77 | 6.7E-04 |
| regulation of cell differentiation | 3.40 | 6.7E-04 |
| regulation of cell migration | 5.16 | 6.7E-04 |
| ***miR-17-5p*** |  |  |
| organelle organization | 1.44 | 2.1E-10 |
| cytosol | 1.33 | 2.4E-10 |
| cytoplasm | 1.15 | 6.4E-09 |
| intracellular organelle | 1.13 | 6.4E-09 |
| regulation of metabolic process | 1.26 | 2.2E-08 |
| intracellular membrane-bounded organelle | 1.15 | 2.7E-08 |
| nucleoplasm | 1.35 | 4.0E-08 |
| enzyme binding | 1.51 | 1.4E-07 |
| protein binding | 1.13 | 1.4E-07 |
| regulation of gene expression | 1.32 | 1.6E-07 |

Based on searching against experimentally-confirmed targets using miRPathDB v2.0.

Selected differentially abundant cord serum miRNAs were validated using qRT-PCR in a different set of biological samples. In the validation sample set, women with obesity again had higher pre-pregnancy BMI and heavier babies and were more frequently of Hispanic ethnicity, but were otherwise similar to women with healthy weight (**Table 5**). In line with the RNASeq data, miR-142-5p and miR-17-5p were more abundant in cord serum from women with obesity compared to women with healthy weight (**Fig. 3B**). MiR-142-3p also tended to be higher in women with obesity, albeit the effect was not statistically significant (**Fig. 3B**). We were unable to validate the downregulated miRNAs identified by RNASeq, and abundances of miR-100-5p and miR-483-5p were similar in samples from the two study groups (P>0.05). Neither miR-149- 3p nor miR-5010-5p were detectable within 40 cycles by qRT-PCR.

**Table 5.** Clinical characteristics of human study participants providing cord serum for qPCR validation of miRNA abundance.

|  | **Healthy weight** | **n** | **Obesity** | **n** | **P value** |
| --- | --- | --- | --- | --- | --- |
| **Pre-pregnancy BMI, kg/m<sup>2</sup>** | 22.5 ± 1.4 | 18 | 36.4 ± 4.6 | 21 | **\*P<0.001** |
| **Ethnicity, number Hispanic (%)** | 0 (0%) | 18 | 11 (52%) | 21 | **\*P<0.001** |
| **Maternal age, years** | 32.0 ± 5.0 | 18 | 30.0 ± 4.7 | 21 | P=0.211 |
| **Gestational age at delivery, weeks** | 39.1 ± 0.7 | 18 | 39.0 ± 1.4 | 21 | P=0.786 |
| **Gestational diabetes, number (%)** | 0 (0%) | 18 | 0 (0%) | 21 | P>0.999 |
| **Vaginal delivery, number (%)** | 2 (11%) | 18 | 0 (0%) | 21 | P=0.207 |
| **Birth weight, kg** | 3.23 ± 0.34 | 17 | 3.72 ± 0.83 | 21 | **\*P=0.019** |
| **Placenta weight, g** | 613 ± 144 | 17 | 734 ± 224 | 19 | P=0.076 |
| **Sex, number female (%)** | 7 (39%) | 18 | 8 (38%) | 21 | P=0.960 |

2 Continuous variables compared by Student's t-test. Categorical variables compared by Chi-squared test. Welch's correction was used if variances 3 differed between groups. Mean ± SD.

### Effect of microRNA mimics on cardiomyocytes *in vitro*

Transfecting NRVMs with synthetic mimics for either miR-142-3p, miR-142-5p or miR-17-5p increased corresponding microRNA expression in the cardiomyocytes (**Fig. 3C-E**). Transfection with the miR-17-5p mimic also increased NRVM expression of miR-142-5p but there was no other crosstalk between the selected microRNAs (**Fig. 3E**). The miR-142-3p mimic reduced *Myh7:Myh6* expression ratio in the transfected cardiomyocytes without affecting *Anf*, *Bnp* or *Serca* expression (**Fig. 3F**), whilst the miR-142-5p mimic did not affect expression of any of the markers of hypertrophy (**Fig. 3G**). MiR-17-5p reduced cardiomyocyte expression of *Anf* and the *Myh7:Myh6* ratio but did not affect *Bnp* or *Serca* (**Fig. 3H**). None of the microRNA mimics altered cardiomyocyte size (miR-142-3p 1297 ± 531 µm^2^, miR-142-5p 1578 ± 829 µm^2^, miR-17-5p 1392 ± 458 µm^2^, P>0.05 in all cases for paired t-test versus SCR 1285 ± 397 µm^2^, n=4 independent cell isolations).

### Effect of primary human trophoblast conditioned medium on cardiomyocytes *in vitro*

Conditioned medium was collected from cultured primary human trophoblast cells isolated from placentas of women with healthy weight or obesity and used to supplement the medium of separately-cultured NRVMs. Women with obesity had higher pre-pregnancy BMI than women with healthy weight but all other clinical characteristics were similar in the two groups (**Table 6**). Trophoblast conditioned medium from placentas of women with obesity increased *Anf* and *Bnp* expression in NRVMs, compared with trophoblast medium from women with healthy weight (**Fig. 4A, B**). By contrast, conditioned medium from trophoblasts of either healthy weight women or women with obesity increased the *Myh7:Myh6* ratio in NRVMs, compared to unconditioned medium (**Fig. 4C**). There was no effect of trophoblast conditioned medium on NRVM *Serca* expression (**Fig. 4D**).

**Figure 4.**
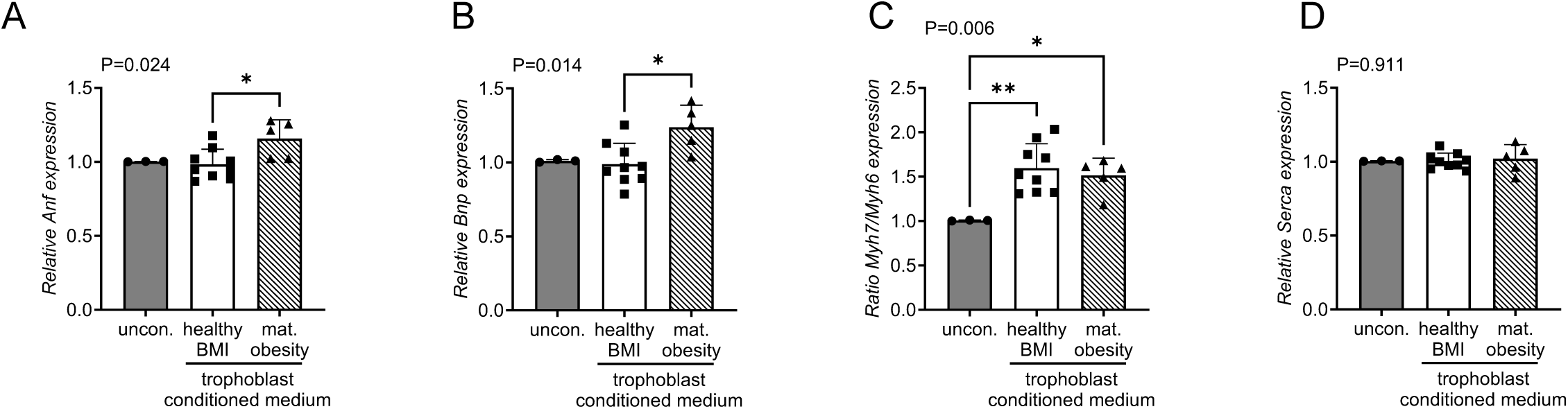
Effect of primary human trophoblast conditioned medium on cardiomyocytes. (A-D) mRNA expression of fetal gene program markers of pathological hypertrophy in NRVMs supplemented with unconditioned medium (n=3) or conditioned medium from primary human trophoblasts isolated from women with healthy weight (con, n=9) or obesity (ob, n=5). Effect of conditioned medium determined by one-way ANOVA with Holm-Sidak post-hoc test. Mean ± SD, points represent individual primary trophoblast cell isolations from different participants.

**Table 6.** Clinical characteristics of human study participants providing primary trophoblast cells.

|  | **Healthy weight** | **n** | **Obesity** | **n** | **P value** |
| --- | --- | --- | --- | --- | --- |
| **Pre-pregnancy BMI, kg/m<sup>2</sup>** | 24.1 ± 2.5 | 7 | 34.2 ± 5.9 | 5 | **\*P=0.002** |
| **Ethnicity, number Hispanic (%)** | 2 (29%) | 7 | 2 (40%) | 5 | P=0.679 |
| **Maternal age, years** | 28.7 ± 6.3 | 7 | 27.6 ± 3.7 | 5 | P=0.732 |
| **Gestational age at delivery, weeks** | 38.9 ± 0.9 | 7 | 38.8 ± 1.1 | 4 | P=0.886 |
| **Gestational diabetes, number (%)** | 0 (0%) | 7 | 0 (0%) | 5 | P>0.999 |
| **Vaginal delivery, number (%)** | 0 (0%) | 7 | 0 (0%) | 5 | P>0.999 |
| **Birth weight, kg** | 3.35 ± 0.38 | 7 | 3.40 ± 0.47 | 5 | P=0.820 |
| **Placenta weight, g** | 648 ± 101 | 7 | 606 ± 93 | 5 | P=0.474 |
| **Sex, number female (%)** | 2 (29%) | 7 | 4 (80%) | 5 | P=0.079 |

6 Continuous variables compared by Student's t-test. Categorical variables compared by Chi-squared test. Welch's correction was used if variances 7 differed between groups. Mean ± SD.

## Discussion

This study shows that circulating factors in human umbilical cord serum from pregnancies complicated by obesity induce hallmarks of pathological hypertrophy in cultured neonatal cardiomyocytes. Cord serum specifically increased neonatal rat ventricular myocyte expression of fetal genes *Anf* and *Bnp* and the ratio of *Myh7:Myh6* expression. Upregulation of these genes was more pronounced when the cardiomyocytes were supplemented with cord serum from fetuses of women with obesity than women with healthy weight. Subjecting the cord serum to

DNase or RNase digestion prevented the effect of maternal obesity on cardiomyocyte gene expression. Maternal obesity increased cord serum abundance of the mature microRNA strands of miR-142 and miR-17, which are associated with cardiac hypertrophy, but miR-142 and miR-17 mimics did not activate the fetal cardiac gene program in cultured neonatal rat ventricular myocytes. Conditioned medium from cultured primary human trophoblast cells from pregnancies complicated by obesity also induced hallmarks of pathological hypertrophy in cardiomyocytes. The findings indicate that fetal circulating microRNAs or placenta-derived factors may mediate cardiomyocyte hypertrophy *in utero* in pregnancies complicated by maternal obesity.

Upregulation of cardiomyocyte natriuretic peptide gene expression and an increase in the *Myh7:Myh6* expression ratio are consistent with the pattern of cardiac gene expression in adults with cardiomyopathy and therefore represent markers of cardiomyocyte dysfunction ^27^. Neonatal rat ventricular myocytes also exhibit these hallmarks when supplemented *in vitro* with human serum from children with dilated cardiomyopathy ^14^. Inotropic factors, such as β- adrenergic agonists, that are mechanistically implicated in cardiac hypertrophic dysfunction cause similar changes in gene expression in cultured cardiomyocytes, alongside increasing cell size ^12^. Our findings therefore indicate that there are circulating factors in human fetuses that have a pathological hypertrophic effect on cardiomyocytes and are more abundant or more potent when pregnancy is complicated by maternal obesity. The transcriptional effect in our study was not accompanied by increased cell size so the change in gene expression is unlikely to be a consequence of, for example, altered intracellular diffusion distance or changes in substrate availability.

Our finding that treating cord serum from pregnancies complicated by obesity with nucleases prevents its effect on cardiomyocytes partly agrees with another study showing that DNase digestion prevents the hypertrophic effect of serum from paediatric dilated cardiomyopathy patients ^28^. However, in contrast with our results, RNase digestion exacerbated, rather than prevented, the hypertrophic effect of paediatric dilated cardiomyopathy in that study ^28^. Our results are more in line with studies in mice with obesity, which show that increased levels of circulating microRNAs originating from adipose tissue and other metabolic organs increase cardiac dysfunction and specifically affect cardiomyocytes ^29,30^ . Circulating microRNAs are stabilised by carrier proteins or packaged in extracellular vesicles. The repeated heat-freeze cycling used to prepare serum samples for our nuclease experiment disrupts these carrier proteins and extracellular vesicles ^31^. Therefore, our findings may indicate that extracellular vesicle or protein-bound microRNAs are pro-hypertrophic factors in cord serum from fetuses of women with obesity.

The increased abundance of extracellular vesicle miR-142 in cord serum from fetuses of women with obesity is consistent with previous studies showing that circulating levels of miR-142-3p are elevated in children ^32^ and adults with obesity ^33,34^. Circulating miR-17-5p is increased in abundance in pregnant women with gestational diabetes ^35^. Maternal circulating microRNA abundance is also altered in pregnant women with obesity, although miR-142 and miR-17 are not amongst the differentially abundant microRNAs identified previously ^17^. Both miR-142-3p and -5p expression is increased in adipose tissue from mice with diet-induced obesity ^36^.

Separately, increased circulating miR-142 and miR-17 abundance is associated with both paediatric and adult cardiomyopathy ^37,38^. MiR-142 and miR-17 control glycolytic and oxidative metabolism in skeletal muscle and tumour cells, respectively ^39,40^. Thus, it is plausible that these specific microRNAs modulate the metabolic switch that occurs normally in fetal cardiomyocytes around birth and cause them to dysfunction when they are upregulated by maternal obesity. Indeed, experimentally induced diabetes in pregnant mice upregulates fetal cardiac miR-142 expression ^41^ .

Despite this, transfecting cardiomyocytes with synthetic miR-142-3p, miR-142-5p and miR-17- 5p mimics did not induce the gene expression hallmarks of pathological hypertrophy in our study. This appears to be in contrast with previous gain-of-function studies showing that *in vivo* miR-142 overexpression exacerbates cardiac dysfunction in adult mice ^25^ and miR-17-3p transfection increases cardiomyocyte size *in vitro* ^26^. These discrepant findings may be explained by differences in the duration or magnitude of the gain-of-function, differing conditions *in vitro* versus *in vivo*, or strand-specific effects of the mature microRNAs. Alternatively, fetal cardiomyocytes may respond differently to these miRNAs than the adult heart. Endogenously expressed microRNAs are likely to act at much lower concentrations than experimentally transfected microRNAs and their *in vivo* cardiac effects are likely to be mediated in combination with other humoral and physical factors. Moreover, loss-of-function studies suggest that endogenous miR-142 and miR-17 are required for more complex roles in physiological cardiac growth ^25,26^. Thus, whilst our findings do not support the concept that miR- 142 or miR-17 directly mediate the pathological effect of cord serum on cardiomyocytes *in vitro* they may still be consistent with a role in modulating fetal cardiac development *in vivo* in pregnancies complicated by obesity.

Our data indicating a pathological hypertrophic effect of the primary trophoblast secretome on cultured cardiomyocytes is in line with previous studies showing that trophoblast stem cells secrete factors that regulate cardiomyocyte differentiation, cardiomyocyte apoptosis and myocardial growth and function ^42–44^. They are also in line with mouse studies showing that placenta-specific gene manipulations lead to alterations in fetal cardiac development ^20^.

Primary human trophoblast cells secrete peptides and extracellular vesicles that contain microRNAs ^22,23,45^. Previous studies indicate that extracellular vesicles mediate communication between trophoblasts and cardiomyocytes ^42,43^. Obesity alters placental microRNA expression and the placenta specific miR profile in the fetal circulation ^16,46^. Therefore, our findings may indicate that altered trophoblast secretion of factors such as microRNAs contributes to the adverse effect of maternal obesity on the fetal heart.

## METHODS

### Study participants

Pregnant women (n=62) with obesity (pre-pregnancy body mass index > 30) or healthy weight (pre-pregnancy body mass index 20-24.9) were recruited to the study prior to delivery at University of Colorado Hospital, with informed consent and ethical approval from the Institutional Review Board of the University of Colorado. Inclusion criteria for both study groups included singleton pregnancy and maternal age 18-45 years. Exclusion criteria included infection, fetal anomaly, maternal smoking, or alcohol or recreational drug use during pregnancy. Umbilical cord blood and/or placenta were collected at delivery and processed within 30 minutes.

### Cord blood and serum handling

Cord blood collected from 55 participants was allowed to coagulate then serum was processed by centrifugation and aliquoted and stored at -80°C. Frozen aliquots of 11 of these cord serum specimens were subjected to rapid heating/re-freezing and nuclease digestion as described ^31^. Briefly, serum was heated to 65°C for 5 minutes then frozen in an ethanol-dry ice bath, in three repeated cycles. This method enhances microRNA detection in human serum samples (Mariner et al, 2016). Subsequently, serum nucleic acids were digested with either DNase (0.1U TURBO DNase per µl serum, AM1907 Thermo Fisher Scientific, Waltham, MA, USA), RNase (750pg RNase A per µl serum, EN0531 Thermo Fisher Scientific, Waltham, MA, USA), or both DNase and RNase consecutively. Enzymes were inactivated after the digest and treated serum was used immediately in subsequent experiments.

### Primary human trophoblast cell isolation and culture

Freshly-dissected placental villous tissue from 12 participants was used for isolation of primary human trophoblast cells, as described ^47^. Briefly, washed and minced tissue was dispersed with trypsin and DNase and cells were subsequently separated by centrifugation over a discontinuous percoll gradient. Trophoblast cells were plated at 5 x 10^6^ cells per dish in 5ml of complete medium (50:50 Ham’s F12:Dulbecco’s Modified Eagle’s Medium, with 10% fetal bovine serum and antibiotics). Culture medium was changed daily thereafter and an aliquot was collected for measurement of human chorionic gonadotrophin, to confirm biochemical differentiation of the cells into syncytiotrophoblast (ELISA from IBL America, Minneapolis, MN, USA). At the final medium change at 66 hours in culture, medium serum content was reduced to 1% and conditioned medium was thereafter collected at 90 hours and stored at -80°C.

### Neonatal rat ventricular myocyte isolation and culture

Primary neonatal rat ventricular myocytes (NRVMs) were isolated from 1- to 2-day-old rat pups as described ^12^. Animal procedures were conducted with approval from the Institutional Animal Care and Use Committee of the University of Colorado Anschutz Medical Campus (#00235).

Briefly, timed-mated pregnant Sprague-Dawley rats (n=19) were purchased from Charles River Laboratories (MA, USA) and allowed to deliver at term. Dams were euthanised by CO_2_ - asphyxiation and cervical dislocation. Pups were killed by decapitation and their hearts immediately excised. Ventricles were dissected, minced and digested in trypsin. Resultant cells underwent an initial short, ∼45-minute plating to remove non-myocytes. Unattached NRVMs were then plated at 3.2 x 10^5^ cells per well on gelatine-coated 6-well plates in Minimum Essential Medium with 5% calf serum and vitamin B12. After 24 hours, NRVMs were transferred to serum-free medium, with insulin, transferrin, bovine serum albumin, vitamin B12 and bromodeoxyuridine, to inhibit cell proliferation. All treatments were subsequently added to the medium and the final outcome measurements were made 72 hours later, without further medium change.

### NRVM treatment

#### Effect of maternal obesity

Cord serum samples from study participants with healthy weight (n=10) and obesity (n=9) were added to serum-free NRVM medium at 2% (v/v) as described ^14^. Control NRVMs remained in serum-free medium. Each serum sample was added to at least 2 technical replicate wells of NRVMs, which were analysed separately then averaged before statistical analysis so that the experimental unit was the human study participant.

#### Effect of serum nucleic acids

Cord serum samples subjected to heat-freeze treatment and originating from participants with both healthy weight and obesity were added to NRVM medium at 2% (v/v). Liposomal transfection reagent (Lipofectamine RNAiMAX, 7.5 µl) was included in the medium to facilitate cellular uptake of serum nucleic acids after the heat-freeze. To determine the effect of depleting serum nucleic acids, heat-freeze treated cord samples were digested with DNase and/or RNase before addition to the medium.

#### Effect of microRNA mimics

NRVMs were transfected with mirVana miRNA mimics for miR-142- 3p, miR-142-5p and miR-17-5p (20µM, **Table 7**) using 7.5µl Lipofectamine RNAiMAX per well.

**Table 7.**
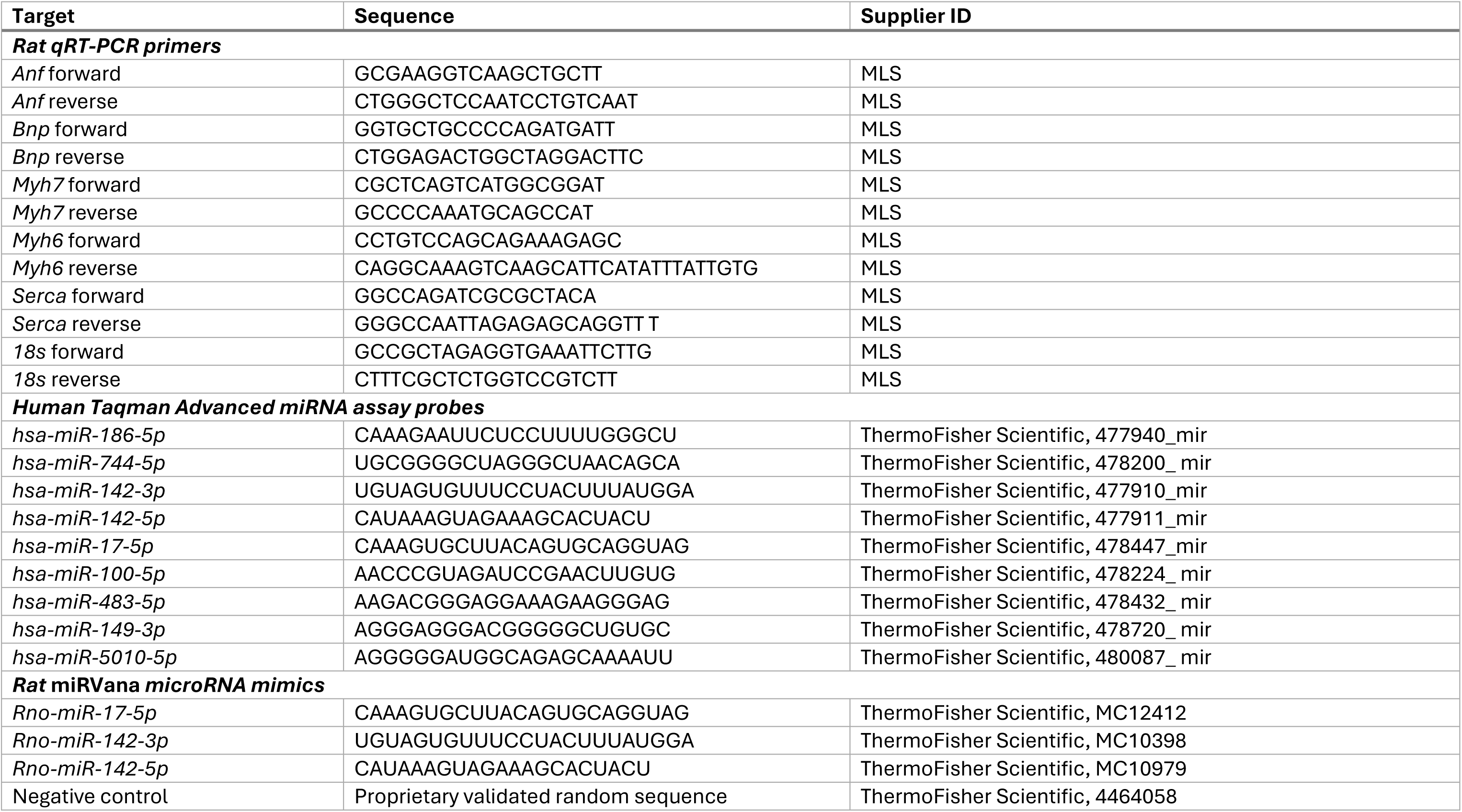
Primer, probe and microRNA mimic sequences.

Transfections were performed in two technical replicate NRVM wells, which were averaged prior to statistical analysis. The experiment was repeated in 3-6 independent biological replicate batches of NRVMs, isolated from primary tissue on different days, so that the experimental unit in the final analysis was the primary cell isolation batch.

#### Effect of primary human trophoblast cell conditioned medium

Conditioned medium from primary human trophoblast cells isolated from participants with healthy weight (n=7) or obesity (n=5) was also added to NRVM medium at 2%. Controls were supplemented with unconditioned trophoblast cell medium.

### Gene expression

NRVMs were lysed and RNA purified 72 hours after treatment, using TRIzol reagent and chloroform extraction. RNA was reverse transcribed (High Capacity cDNA Reverse Trascription Kit, Applied Biosystems) then expression of the rat mRNAs *Anf, Bnp, Myh7, Myh6* and *Serca* was determined using specific primers and SYBR green chemistry. Expression of *Anf* and *Bnp* and the ratio *Myh7:Myh6* are commonly higher in both fetal hearts and failing adult hearts than in healthy adult hearts so they are collectively called the fetal gene program and used as readouts of pathological cardiomyocyte hypertrophy ^27^. Expression was determined relative to *18s* mRNA, using the ddCt method. Primer sequences are given in **Table 7**.

### Immunostaining and cell size measurement

NRVMs were either visualised live and unstained or fixed in 4% paraformaldehyde then immunostained using an anti-α-actinin primary antibody (1:2000 overnight) and Alexa-fluor 488- conjugated anti-mouse secondary antibody (1:1000, 45 minutes). Two-dimensional images were acquired using an EVOS FL Auto Imaging System (Thermo Fisher Scientific). Cell area was then measured using Image J.

### Isolation of RNA from cord serum extracellular vesicles

Total RNA was isolated from extracellular vesicles in 51 cord serum specimens using a commercially available kit (exoRNeasy Serum/Plasma, Qiagen, Germantown, MD, USA). Deep frozen cord serum was thawed by brief incubation at 37°C then filtered using a 0.8µm syringe filter. Extracellular vesicles were isolated from 400µl of filtered serum using membrane-based affinity binding spin columns. Vesicles were lysed and RNA isolated using a phenol/guanidine-based approach with chloroform extraction, ethanol precipitation and silica membrane purification. A synthetic miRNA (5.6 x 10^8^ copies of C. elegans miR-39) was spiked into the vesicle lysate, prior to chloroform extraction, as an internal control for RNA recovery in the isolation procedure.

### Small RNASeq

Small RNA libraries were prepared from 6µl total extracellular vesicle RNA (Small RNA-Seq Library Prep Kit, Lexogen, NH, USA). Small RNASeq was then performed with 150 cycles of paired-end reads using the Illumina NovaSEQ 6000, in collaboration with the Genomics and Microarray Core facility of the University of Colorado Anschutz Medical Campus. Resultant reads were trimmed of adapter sequences using BBDuk (https://jgi.doe.gov/data-and-tools/software-tools/bbtools/bb-tools-user-guide/bbduk-guide/) with only reads between 15-31 bases kept. The reads were aligned to miRbase human mature miRNA sequences ^48^ using Burrows-Wheeler Aligner ^49^. Counts were compiled and converted to counts per million. Low abundance microRNAs were removed by requiring greater that 20 counts in at least 12 samples. Differentially expressed microRNAs was calculated using EdgeR ^50^. Gene ontology pathways significantly enriched with mRNA targets of the differentially expressed microRNAs were identified by filtering for experimentally-confirmed interactions using miRPathDB 2.0 ^51^.

### microRNA qPCR

Differentially expressed miRNAs in cord serum extracellular vesicles were validated in a separate set of biological samples from participants with obesity or healthy weight, using primer/probe-based Taqman Advanced miRNA Assays and the Taqman Advanced miRNA cDNA Synthesis Kit (both Life Technologies Corp., CA, USA). Using 2µl of total extracellular vesicle RNA, mature miRNAs were extended with 3’ poly(A) addition and 5’ adaptor ligation then reverse transcribed and uniformly amplified. Relative miRNA quantity was determined by qRT-PCR, using the ddCt method normalised to the geometric mean of endogenous (human miR-186-3p and miR-744-5p) and exogenous (spiked-in) control sequences (C. elegans miR-39). Taqman assay IDs are given in **Table 7**.

### Statistics

Results are presented as mean ± SD. Clinical characteristics and cord serum microRNA abundances were compared between participants with healthy weight and obesity by Student’s t-test for continuous variables or Chi-squared test for categorical variables. Effects of serum treatments, microRNA mimics and conditioned medium on cardiomyocytes were determined by Student’s t-test or one-way ANOVA with Sidak post-hoc test. Interacting effects of maternal obesity and serum heat-freezing/digest were determined by mixed effects model. Homogeneity of variances was assessed by F-test and Welch’s correction was used if variances differed between groups. Distribution was assessed by Shapiro-Wilk test and data were log transformed if they were not normally distributed. Effects were considered statistically significant when P<0.05.

## CONCLUSION

Overall, the results support our hypothesis by showing that circulating factors in serum from fetuses of women with obesity induce hallmarks of pathological hypertrophy in neonatal cardiomyocytes. They also identify fetal circulating microRNAs that are altered in abundance with maternal obesity and indicate that pro-hypertrophic fetal circulating factors could originate from the placenta. Our findings imply that fetal circulating microRNAs and placenta-derived factors contribute mechanistically to the adverse effects of maternal obesity on fetal development and offspring cardiometabolic disease risk. We speculate that interventions targeting circulating microRNAs or altering placental function could prevent offspring cardiac dysfunction in pregnancies complicated by maternal obesity.

## Acknowledgements

This study was supported by the Ludeman Family Center for Women’s Health Research and National Institutes of Health R01HD065007, P30DK048520 via Colorado Nutrition Obesity Research Center Pilot and Feasibility Program, and P30CA046934 via the University of Colorado Cancer Center Bioinformatics and Biostatistics Shared Resource.

